# Physical Chemistry of Drug Permeation through the Cell Membrane with Atomistic Detail

**DOI:** 10.1101/2023.07.24.550356

**Authors:** Mirko Paulikat, GiovanniMaria Piccini, Emiliano Ippoliti, Giulia Rossetti, Fabio Arnesano, Paolo Carloni

## Abstract

We provide a molecular-level description of the thermodynamics and mechanistic aspects of drug permeation through the cell membrane. As a case study, we considered the anti-malaria, FDA approved drug chloroquine. Molecular dynamics simulations of the molecule (in its neutral and protonated form) were performed in the presence of different lipid bilayers, with the aim of uncovering key aspects of the permeation process, a fundamental step for drug’s action. Free energy values obtained by well-tempered metadynamics simulations suggest that the neutral form is the only permeating protomer, consistent with experimental data. H-bond interactions of the drug with water molecules and membrane headgroups play a crucial role for permeation. The presence of the transmembrane potential, investigated here for the first time in a drug permeation study, does not qualitatively affect these conclusions.

**TOC Graphic:** 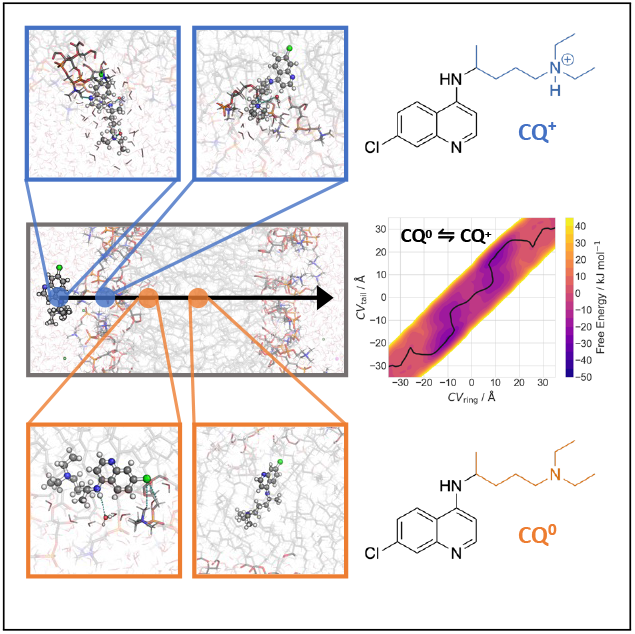

---

The interaction of drugs with biological membranes impact dramatically on their mechanism of action. A typical example is the FDA-approved antimalarial drug chloroquine (**CQ**).^1, 2^ This drug also possesses antiviral,^3–5^ antirheumatic,^6–8^ and anti-inflammatory properties.^9, 10^ In fact, its beneficial effect stems from the drug’s ability to reach the food vacuole, an acidic compartment of the parasite, permeating several biological membranes.^11^ The drug can feature three different protonation species at the extracellular pH (Chart 1), from the neutral one (**CQ**^0^), to the mono-(**CQ^+^**) and di-protonated (**CQ^2+^**) species (p*K*_a1_ and p*K*_a2_ are 8.1 and 10.4 at 310 K).^12^

**Chart 1:**
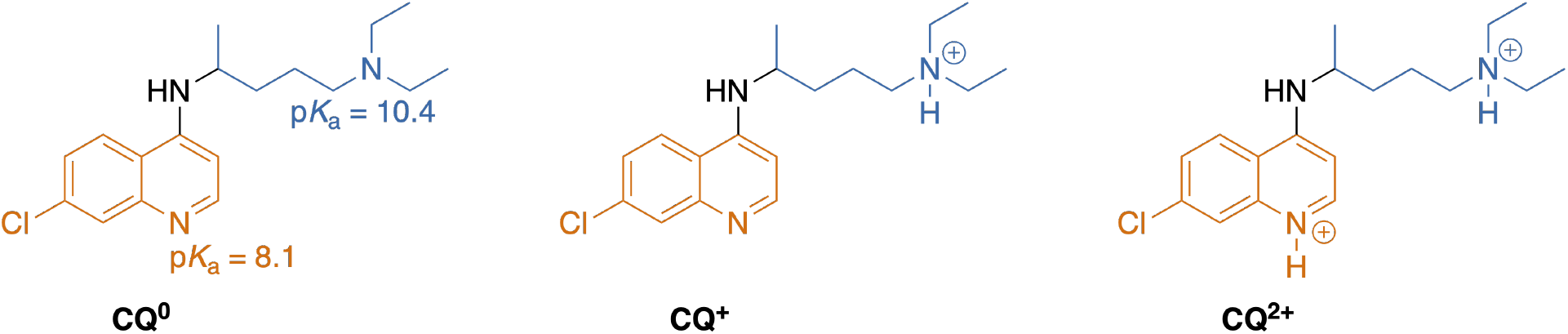
Protomers of chloroquine (**CQ**). The ring and tail moieties are colored in orange and blue, respectively.

**CQ** penetrates malaria-infected human erythrocytes by passive diffusion of the free base, which may become protonated and trapped inside the acidic vacuole. ^13^ However, a detailed study of **CQ** binding to intact and lysed erythrocytes indicated that the mechanism of **CQ** accumulation in intact cells is actually a combination of ion trapping at acidic pH (a consequence of the basic nature of the drug and the pH gradient across the membrane) and binding to cell components.^14^

The analysis of **CQ** uptake into erythrocytes infected with drug-sensitive and resistant strains of the human malaria parasite *Plasmodium falciparum* revealed that uptake can be resolved into a non-saturable and a saturable component.^15, 16;1^ Non-saturable uptake in the submicromolar range^17^ is nonspecific and is attributed to low-affinity binding of **CQ** to plentiful cytosolic proteins.^18^ Conversely, saturable uptake at nanomolar drug concentrations is important for antimalarial activity^17^ and attributed to intracellular binding of **CQ** to ferriprotoporphyrin IX (FPIX), a product of parasite haemoglobin digestion.^15, 19^ FPIX is polymerized into an inert crystalline substance called hemozoin, but **CQ** inhibits this process causing a buildup of free FPIX and/or **CQ**–FPIX complex that will ultimately kill the parasite.^20–22^ Thus, the intracellular uptake of **CQ** in malaria-infected cells is primarily due to passive diffusion followed by saturable binding of **CQ** to FPIX rather than active import by membrane transporters.^15, 23^

The drugs’ uptake requires about 1 hour. ^24, 25;2^ Permeation occurs by fast passive diffusion with a permeability coefficient of 7.2 cm s^−1^ for **CQ**^0^ at 310 K.^11, 14, 26^ Only **CQ**^0^ permeates the membrane in spite of being extremely scarce in physiological conditions (about one part for 10,000).^14, 26^ While one can provide simple electrostatic arguments to explain this – charged molecules are not thermodynamically stable inside the hydrophobic part of the membrane – a molecular view of drug permeation, and in particular of the role of water for drug permeation, is still missing.

For the last decade, a variety of state-of-the-art enhanced sampling methods have successfully described the structure, dynamics and energetics of small molecule permeation.^27–33^ Here we use well-tempered metadynamics (WTMetaD), an exact method to calculate the free energies of a process as a function of appropriate collective variables,^34, 35^ to describe the process. Our calculations are carried out for the two species **CQ**^0^ and **CQ^+^**, in model membranes with different compositions or in the presence of a transmembrane potential. **CQ**^0^ (and not **CQ^+^**) passes the membrane, consistent with experimental evidence. The process is not qualitatively changed in the presence of the membrane potential.

As a first step, the free energy associated with the permeation of the **CQ** molecule is calculated using well-tempered metadynamics, which is an exact method to calculate the free energy landscapes as a function of apt CVs.^35^ Previous studies have used either one or two CVs.^31, 33, 36^ Choosing an additional CV has shown to significantly improve the results.^31^ Here, we choose the following CVs: the distances between the center of the membrane and the ring/tail moieties of **CQ** (Figure 1).

**Figure 1:**
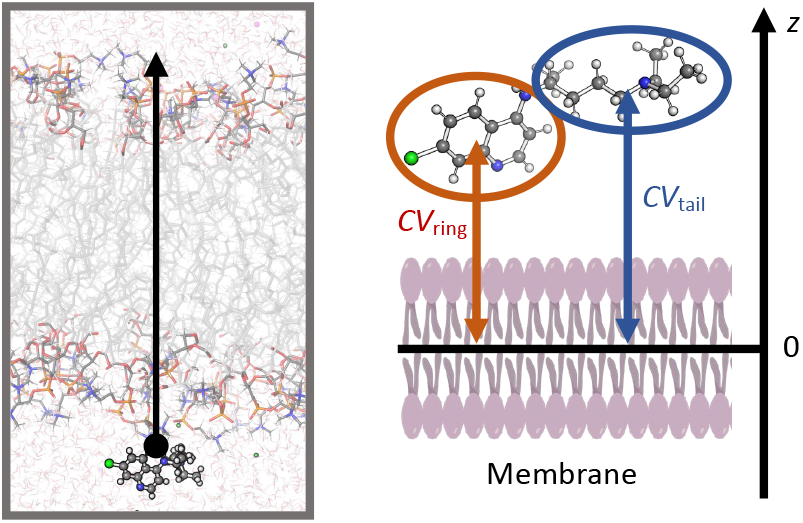
Permeation coordinate. Left: Snapshot of chloroquine in the presence of a lipid bilayer, schematically indicating the permeation coordinate with the black arrow. Right: The two collective variables, CV_ring_ and CV_tail_, measure the distances of the ring/tail moieties of chloroquine with respect to the membrane center along the normal *z*-axis.

## **Permeation of CQ**^0^

MD simulations of **CQ**^0^ (Chart 1) in explicit solvent and in the presence of the POPC membrane (Figures 1–2) show that all H-bond functionalities of **CQ**^0^ interact with the solvent (Figure S1). After leaving the bulk solvent, the ring and tail moieties of **CQ**^0^ interact with POPC headgroups, while the rest is still fully solvated (Figure 2a). Then, the ring moiety dives into the membrane (Figure 2b) and the tail interacts, in turn, with the POPC headgroups (Figure 2b). In this step, the free energy decreases by 20 kJ mol*^−^*^1^ (Figure 2, left). Subsequently, **CQ**^0^ forms H-bonds with POPC headgroups and water molecules, while the remainder interacts with the hydrophobic core of the membrane (Figure 2c). This is the global free energy minimum (*−*40 kJ mol*^−^*^1^ lower than in the solvated state), located inside the membrane. Then, the free energy increases by 11 kJ mol*^−^*^1^: **CQ**^0^ is entirely inside the membrane, interacting exclusively with the hydrophobic core (Figures 2d and 2e). The second part of the translocation is, as expected, completely symmetrical with respect to the first part (Figure 2f). We conclude that **CQ**^0^ is located at the water/membrane interfaces, so as to form H-bond interactions with its polar moiety and hydrophobic with the rest. It can permeate from side to side in the sub-ms timescale, taking into account the barrier of about 10 kJ mol*^−^*^1^. This is consistent with experimental evidence, which shows that the molecule can permeate through the membrane.

**Figure 2:**
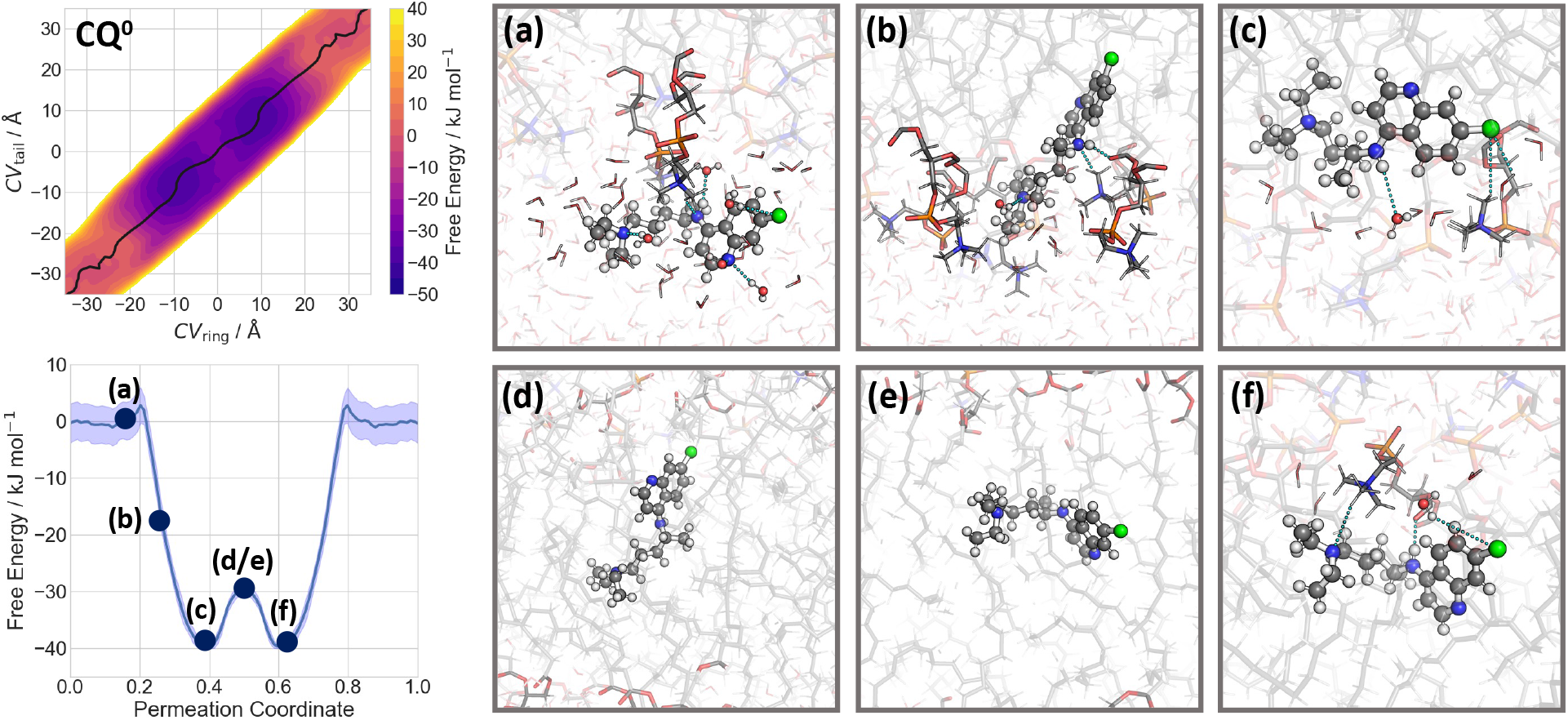
Permeation of **CQ**^0^ through a POPC membrane. Left: the free energy of the process, calculated as a function of CV_ring_ and CV_tail_ (see Figure 1). The minimum free energy path is plotted as black line (top) and is shown in its one-dimensional representation below. (a)–(f) Representative snapshots of the **CQ**^0^ permeation process across the POPC membrane. The associated free energies are indicated in the minimum free energy profile.

## Protonated species

The molecular mechanism of permeation of **CQ^+^** is similar to that of **CQ**^0^. However, in this case the global minimum is far more solvated than in the case of the neutral molecule, most likely because the solvent stabilizes the charged molecule (Figure 3b). Here, the ring moiety interacts with the hydrophobic core of the membrane, while the charged tail of **CQ^+^** interacts with water molecules and POPC headgroups at the lipid/water interface. More importantly, the maximum free energy inside the membrane now shifts by as much as 35 kJ mol*^−^*^1^ with respect to the solvent state. This is caused by the destabilization of the charged molecule within the membrane along with membrane deformation and water defects (Figure 3c–f). The total energy barrier for membrane translocation is 60 kJ mol*^−^*^1^. Thus, we conclude that, consistent with experiment, the protonated **CQ^+^** does not cross the membrane.

**Figure 3:**
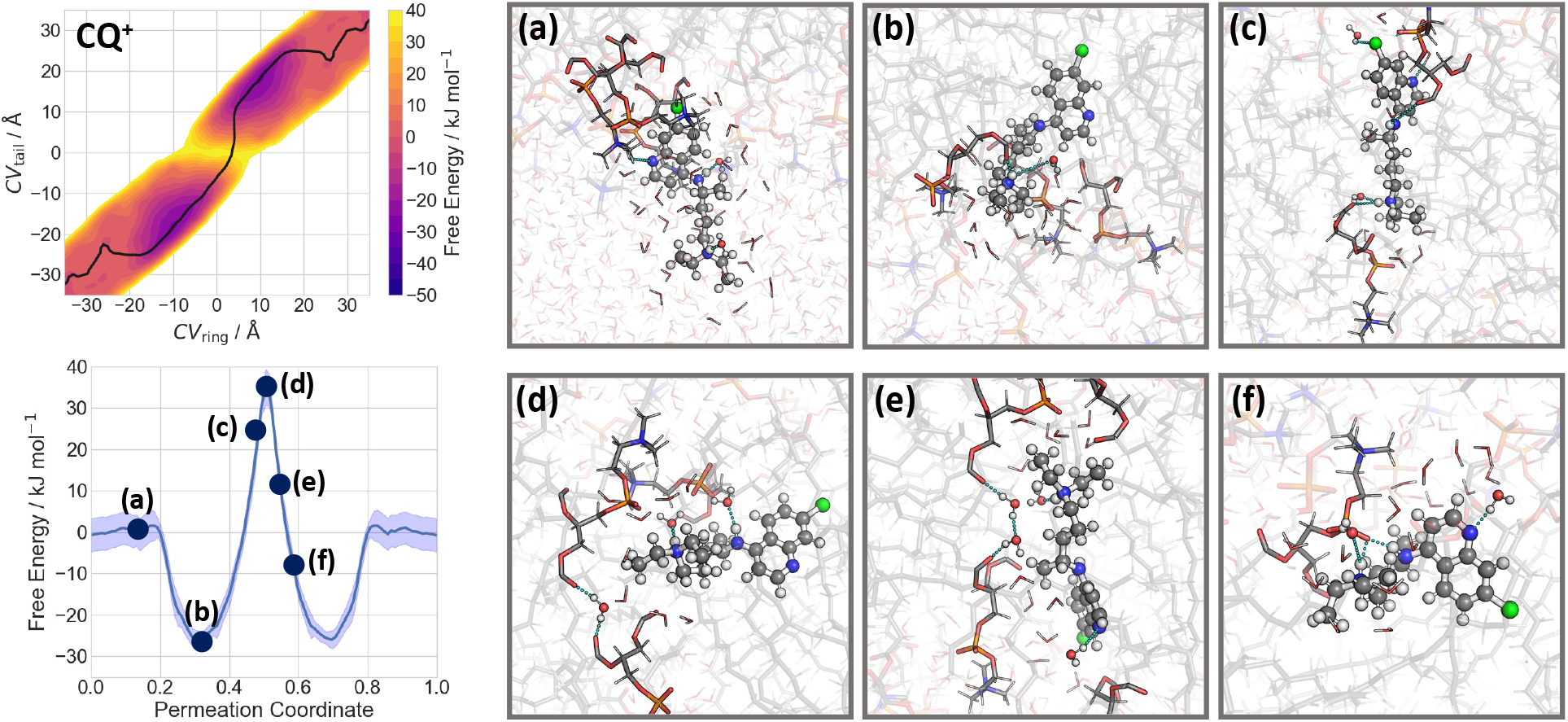
Permeation of **CQ^+^** through a POPC membrane. Left: the free energy of the process, calculated as a function of CV_ring_ and CV_tail_ (see Figure 1). The minimum free energy path is plotted as black line (top) and is shown in its one-dimensional representation below. (a)–(f) Representative snapshots of the **CQ^+^** permeation process across the POPC membrane. The associated free energies are indicated in the minimum free energy profile.

The calculated free energy difference between **CQ**^0^ and **CQ^+^** suggests that the protonated species is not stable within the membrane (Figure 4a). The reversal point is located at the lipid/water interface, when the charged tail of **CQ^+^** enters the hydrophobic core of the membrane (CV_tail_*≈ ±*10 Å). The permeation mechanism can be described as follows: the protonated **CQ^+^** dives into the membrane with its ring moiety, while the tail moiety is still sufficiently solvated at the lipid/water interface (Figures 3b and 4). As discussed above, this corresponds to the global minimum of **CQ^+^**. When the tail moiety enters the hydrophobic core of the membrane, the proton is expected to be transferred to the solution phase. Both water molecules and POPC headgroups interact with the tail moiety and could act as proton acceptors (Figure 3b). The neutral **CQ**^0^ is then located inside the membrane (Figure 2c) and is able to pass the membrane center. The total barrier for membrane translocation is 16 kJ mol*^−^*^1^, consistent with the experimental evidence that **CQ**^0^ can pass the membrane. Upon leaving the membrane, the amine nitrogen gets re-protonated so that the tail moiety exits the hydrophobic core of the membrane first, followed by the ring moiety to complete the permeation process.

**Figure 4:**
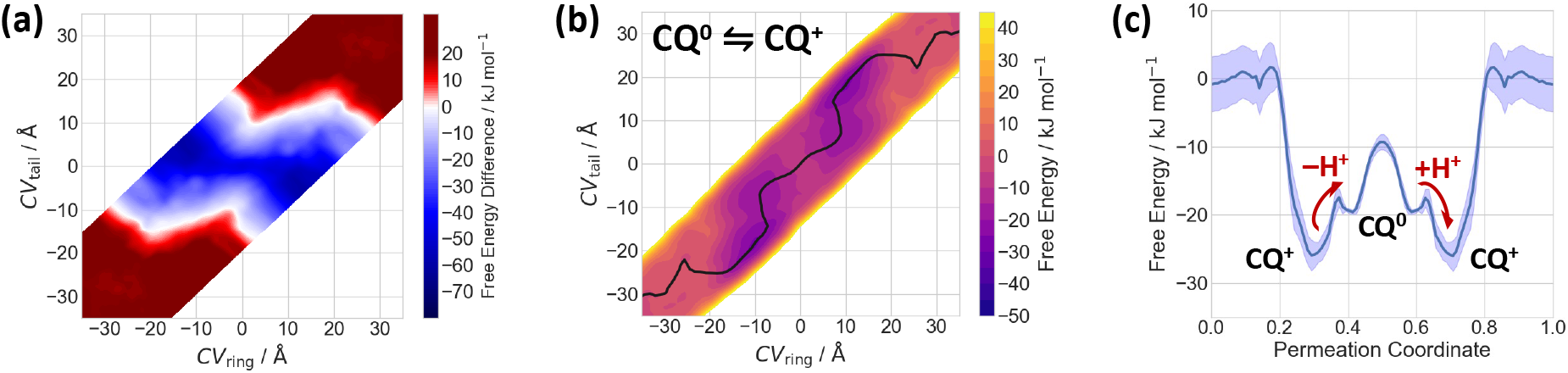
Protonation-dependent permeation of **CQ** through a POPC membrane. (a) The free energy difference of **CQ^+^** and **CQ**^0^, as a function of CV_ring_ and CV_tail_ (see Figure 1). The calculated profile of **CQ**^0^ is shifted with respect to the aqueous state, considering a solution pH of 7 and a p*K*_a_ value of 10.4 at 310 K. Red indicates a preferred protonated state (**CQ^+^**), while blue indicates a preferred neutral state (**CQ**^0^). (b) Free energy surface for the permeation of **CQ**^0^ **⇌ CQ^+^**, as a function of the two CVs, weighted according to the calculated population of the corresponding protonation state. The minimum free energy path is shown as black line. (c) 1D minimum free energy path of the **CQ**^0^ **⇌ CQ^+^** free energy surface. Suggested positions for proton transfer are indicated as red arrows.

In a recent study, the permeation process of a weak base, propranolol, was investigated by MD simulations at constant pH, allowing for dynamic protonation of the molecule in the inhomogeneous environments.^37^ Similar to our findings, the change in protonation state was observed at the lipid/water interface and the free energy profile proceeds along the energetically favorable protomer in the different phases.^37^ We conclude that the protonation states of **CQ** (or similar drug-like molecules) can have a significant impact on the permeation mechanism and energetics even if only the neutral molecule is able to cross the membrane center.

## Impact of membrane composition on permeation

The mechanism of permeation across the POPC/POPS mixed bilayer is similar to that of the pure POPC membrane. However, quantitatively, the energetics of the permeation process differs slightly in the headgroup regions (Figure 5). After leaving the bulk solvent, both the ring and tail moieties of **CQ**^0^ interact with the POPC/POPS headgroups, while the remaining part of the molecule is still fully solvated (Figures S10a and S11a). The free energy increases slightly by 7 kJ mol*^−^*^1^. The increase in free energy is more pronounced than for the pure POPC membrane, possibly due to the disruption of the rather strong interactions between the POPS headgroups (which are negatively charged) and water molecules. Then, the ring moiety dives into the membrane and the other moiety interacts, in turn, with POPC/POPS headgroups (Figures S10b and S11b). The free energy decreases by 20 kJ mol*^−^*^1^. Next, the drug is inside the membrane: the molecule forms H-bonds with POPC/POPS headgroups and water molecules, while the rest interacts with the hydrophobic core of the membrane (Figures S10c and S11c). This is the global free energy minimum (*−*42 kJ mol*^−^*^1^ lower than in the solvated state). Then, the free energy increases by 12 kJ mol*^−^*^1^ when the molecule is completely inside the membrane, interacting only with the hydrophobic core (Figures S10d and S11d). The second part of the translocation is, as expected, symmetrical with respect to the first part. Thus, our simulations suggest that the drug is located at the water/membrane interfaces, so as to form H-bond interactions with its polar moiety and hydrophobic with the rest.

**Figure 5:**
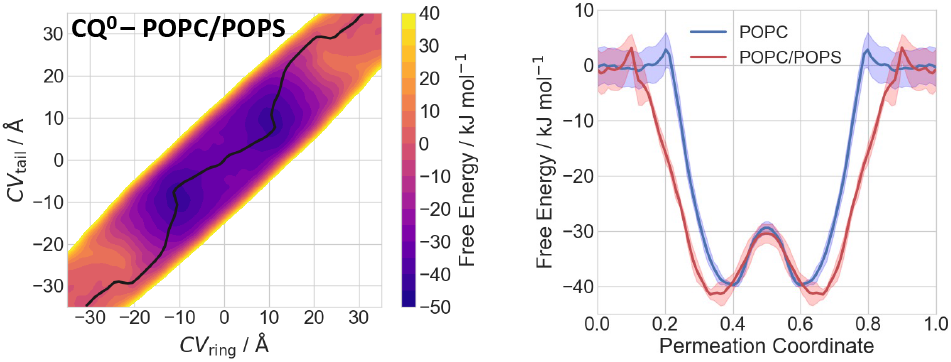
Effect of the membrane composition on the permeation of **CQ**^0^. Left: Free energy surface for the permeation of **CQ**^0^ through a POPC/POPS membrane, as a function of the CV_ring_ and CV_tail_. The minimum free energy path is shown as black line. Right: 1D minimum free energy path of the free energy surface for **CQ**^0^ permeating POPC (blue) and mixed POPC/POPS (red).

**Figure 6:**
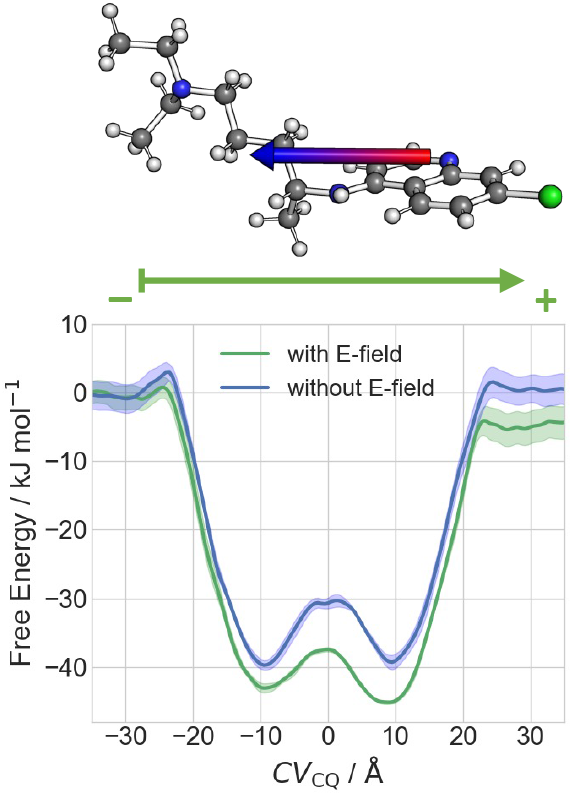
Effect of the external potential on permeation. Top: Ball-and-stick representation of **CQ**^0^ showing its dipole moment as an arrow from negative (red) to positive (blue) charge distribution. Bottom: Comparison of free energies with and without an external field for **CQ**^0^ permeation through the POPC membrane. The direction of the external field is shown as an arrow.

## Impact of external potential on permeation

The effect of the membrane potential on permeation has not been investigated so far, to the best of our knowledge. Here, we compare the energetics of the permeation of **CQ**^0^ with and without an external electric field (E) of 10 mV Å*^−^*^1^ along the POPC membrane. Our field value is not too dissimilar to that widely used in computational electrophysiology setups.^38–41^ To a first approximation, we expect changes in the potential energy of the process of the order of the 2 *E · µ* (*≈* 2 *−* 3 kJ mol*^−^*^1^), where *µ* = 5.8 D is used for the dipole moment of **CQ**^0^. Therefore, even fields much larger than the physiological ones (*≈* 0.5 *−* 3 mV Å*^−^*^1^)^42^ are unlikely to drastically affect permeation. The free energy calculations of **CQ**^0^ permeation using CV_CQ_^3^ show that this is indeed the case. The first free energy minimum is *−*43 kJ mol*^−^*^1^ lower than the value in the solvated state, and 3 kJ mol*^−^*^1^ lower than the value without the external electric field. Thus, **CQ**^0^ slightly prefers the lipid phase when an E-field is taken into account. The free energy increases by 7 kJ mol*^−^*^1^ when the molecule is completely inside the membrane. The second free energy minimum is even lower (*−*46 kJ mol*^−^*^1^). This asymmetry arises from the preferred orientation of **CQ**^0^ along the direction of E (Figure S12). We conclude that the effect of the external electric field is of minor importance for **CQ**^0^ permeation under physiological conditions.

## Permeability Coefficients

As a final step, we calculated the permeability coefficient from the inhomogeneous solubility-diffusion model. The position-dependent diffusion coefficients are given in the SI. The permeability coefficients of **CQ**^0^ are 38.2*±* 7.8 cm s^−1^ and 44*±* 12 cm s^−1^ for the permeation through the POPC and the POPC/POPS model membranes, respectively. Considering the protonation-dependent permeation process (**CQ**^0^ ⇌ **CQ^+^**), the permeability coefficient amounts to 26.0 *±* 6.0 cm s^−1^. The difference between the experimental value (7.2 cm s^−1^ at 310 K for the human erythrocyte membrane)^11, 14^ can be ascribed, at least in part, to the different chemical environments: the real cell membrane differs from our model membranes in several aspects. Most importantly, it features a high content of cholesterol that can significantly affect membrane permeability, usually lowering its value.^43, 44^ In addition, our models also lack sphingolipids that are charged groups and that usually increase the permeability.^45^ So, in the end the membrane permeability will be modified by these chemicals in a highly non-trivial manner.

In conclusion, we present a study of chloroquine permeability along permeation through model membranes. The calculated permeability of the drug is about three to six times higher than experimentally found in biological membranes. Previous drug permeation studies predicted permeabilities with errors similar to that found here. ^27, 28, 31, 33^ The difference might be ascribed, at least in part, by the highly diverse environments and by possible changes due to dynamic protonation of the drug, here not included. ^37^ In addition, we show that both the neutral (**CQ**^0^) and the protonated (**CQ^+^**) species are partially solvated in their global free energy minima at the membrane/water interface. However, while **CQ**^0^ can cross the hydrophobic core of the membrane in the absence of any H-bond interactions, **CQ^+^** requires these interactions at its charged moiety in the permeation process. Consequently, **CQ**^0^ is the only species able to cross the membrane in a time scale compatible with experiments. The impact of cell membrane potential is negligible.

## Computational Methods

All calculations and most of the analyses were carried out with the GROMACS 2019.4 package interfaced with the PLUMED-2.5.3 plugin.^46–48^

### Lipid bilayer preparation

A 2*×*37 pure POPC^4^ bilayer (**M1**) and a 2*×*40 POPC/POPS^5^ (**M2**) bilayer with 7:3 stoichiometry were generated using the CHARMM GUI web server^49^ and inserted in a box of sizes 50 Å *×* 50 Å *×* 100 Å. A 30 Å thick water slabs were located on top of each leaflet. 11 (35) Na^+^ and 11 (11) Cl^–^ ions were added to **M1** (**M2**) so as to ensure electroneutrality and to keep the salt concentration of 150 mM, a value not too dissimilar from the extracellular fluid.^42^ Overlapping water molecules were removed. **M1** and **M2** contained 23,423 and 24,752 atoms, respectively.

### Molecular dynamics parameters

The AMBER Lipid 17,^50^ TIP3P,^51^ Joung-Cheatham^52^ force fields were used for the membranes, water molecules and Na^+^/Cl^–^ ions, respectively. As for the drugs, the GAFF2 force field was employed for bonded and van der Waals parameters of the drugs,^53^ while the atomic partial charges were calculated using the RESP fit method at the HF/6-31G*//B3LYP/6-31G* level of theory.^54–56^ Periodic boundary conditions were applied. Electrostatic interactions were calculated using the particle-mesh Ewald summation method with a real space cutoff of 10 Å.^57^ Lennard-Jones interactions were truncated at the same cutoff value. An analytical correction for the potential energy and for the overall pressure taking into account the truncation of the Lennard-Jones interactions (implemented in GROMACS^46^) has been applied. All bonds involving hydrogen atoms were constrained using the LINCS algorithm.^58^ An integration time step of 2 fs was used. The center of mass motion was removed separately for the lipids and aqueous phase every 100 steps. Unless differently stated, (i) constant temperature simulations were achieved by coupling the systems with a Nośe-Hoover thermostat, using a time constant of 0.5 ps.;^59, 60^ (ii) constant pressure simulations were obtained using semi isotropic Parrinello-Rahman barostat at 1 bar, using  a time constant of 1.0 ps and compressibility of 4.5 10*^−^*^5^ bar*^−^*^1^.^61^

### Molecular dynamics simulation of M1 and M2

The systems were first energy-minimized by 10,000 steepest descent steps. Then, they were heated up by 100 ps from 0 to 310 K by molecular dynamics with velocity rescale methods with a time constant of 0.1 ps.^62^ Here, the lipid atoms were restrained to their position by harmonic restraints of 50 kJ mol*^−^*^1^ Å*^−^*^2^. The systems were then unconstrained. Next, **M1** and **M2** underwent 60 and 110 ns *images/550356v1_Nimages/550356v1_images/550356v1_Pimages/550356v1_images/550356v1_Timages/550356v1_* MD simulations, respectively.

### Insertion of the drugs

The drugs were placed into the aqueous phase at 35 Å along the membrane normal (*z*-axis in Figure 1) from the center of the membrane. We used the GROMACS insert-molecule module.^46^ Also in this case, overlapping water molecules were removed. The resulting systems underwent 10 ns-long *images/550356v1_Nimages/550356v1_images/550356v1_Pimages/550356v1_images/550356v1_Timages/550356v1_* simulations. The final configurations were used for subsequent calculations of the free energy. Selected configurations in which the ligand was located at around *−*40 Å from the center of the membrane along the *z*-axis (*z* = 0) were used for subsequent calculations of the diffusion coefficient. For the calculation of the diffusion coefficient, also a total of 1.275 µs *images/550356v1_Nimages/550356v1_images/550356v1_Vimages/550356v1_images/550356v1_Eimages/550356v1_* trajectories were collected (see Section S6 in the SI).

### Free energy calculations

The free energy as a function of a specific collective variable was calculated by well-tempered metadynamics.^34^ Our collective variables (CVs) are the distances between the drug’s center of masses (COMs) and the COM of 40 selected membrane atoms, representing the center of the membrane. They correspond to the *z*-component of its direction vector (see Figure 1 which shows the *z*-axis). The center of the membrane is set to have *z* = 0. This CV has been used also in previous drug permeation studies: ^27–33^ Gaussians with 1.2 kJ mol*^−^*^1^ initial height and 0.5 Å sigma values were deposited every 2 ps, similarly to Refs. [ 31,36]. The bias factor for **CQ^+^** (25) is larger than that for **CQ**^0^ (20), because a higher free energy barrier of the charged species is expected. The bias potential was evaluated on a grid of spacing of 0.01 Å. The reweighting factor *c*(*t*) was calculated on the fly. Free energy profiles were constructed through the reweighting scheme of Ref. [63].

4.5 µs were simulated for each system. The first 500 ns were not included in the analysis as in Ref. [ 63]. Errors of the free energy profiles were estimated by block averaging with three blocks of the processed data (see SI). The free energy profiles were then symmetrized and checked for their asymmetry (see SI). The minimum free energy paths on the 2D free energy surface were determined by the string method.^64^

The calculations with the electric field were carried out using the GROMACS routine ^46^ to mimic the presence of a transmembrane potential. A potential of 10 mV Å*^−^*^1^ was set. Not too dissimilar potentials have been used in other setups, such as the computational electrophysiology simulations.^38–41^ 3 µs were simulated with and without the electric field, using a simpler one-dimensional CV space (see Figure S12a in SI).

### Calculated properties

i. The diffusion and the permeability coefficient were calculated as described in Section S6 of SI.^65, 66^
ii. Radial distribution functions of the water-drug H-bonds were calculated from the well-tempered metadynamics runs for **M1**. All snapshots in which the drug was located in the aqueous phase with a distance larger than 30 Å from the membrane center (10,403 and 14,440 structures for **CQ**^0^ and **CQ^+^**, respectively) were used.
iii. The free energy differences between **CQ**^0^ and **CQ^+^** were calculated as in Refs. [30,67].

The free energy of **CQ**^0^ is first shifted to account for its acid-base equilibrium in the aqueous phase, considering p*K*_a2_ (10.4) at the conditions of the simulation (neutral pH and temperature of 310 K).^6^ The **CQ**^0^ ⇌ **CQ^+^** free energy surface is then determined by weighting the individual free energy surfaces with respect to the calculated population of protomers as a function of CVs (see SI - Section S3 for details). (iv) The dipole moment of **CQ**^0^ was calculated at the B3LYP/6-31G* level of theory using the Gaussian09 code.^68^

## Acknowledgement

The authors gratefully acknowledge the Gauss Centre for Supercomputing e.V. (www.gauss-centre.eu) and the Jülich Supercomputer Center for providing the computing time for this project. We also thank the University of Bari and the Italian Ministero dell’Università e della Ricerca (PRIN 2017WBZFHL to F.A.) for support. GR and PC acknowledge the Helmholtz European Partnering fundings for the project ‘Innovative high-performance computing approaches for molecular neuromedicine’, as well as the Joint Lab ‘Supercomputing and Modeling for the Human Brain’ of the Helmholtz Association, Germany. GR also acknowledges the Federal Ministry of Education and Research (BMBF) and the state of North Rhine-Westphalia as part of the NHR Program.

## Supporting Information Available

Hydrogen bond analysis of **CQ** in the aqueous phase; detailed analysis of free energy surfaces; representative snapshots showing the permeation across the membrane models; description of the calculation of the **CQ**^0^ ⇌ **CQ^+^** free energy surface; computational details on the calculation of the diffusion coefficients and permeability coefficients (PDF).

## Footnotes

(1 The uptake is different for healthy erythrocytes. Indeed, the saturable high-affinity component of **CQ** (i.e., FPIX) is absent or deficient in uninfected cells due to lack of hemoglobin degradation by the parasite.^17^ As a matter of fact, there is no evidence of saturation of **CQ** transport in uninfected erythrocytes. ^14^

(2 Using tritiated or fluorescently-tagged **CQ**, at least two phases were observed: an extremely rapid short phase (*<* 30 s), followed by a slower phase leading to steady state within 1 hour. ^24, 25^

(3 In these simulations, we used a simplified one-dimensional CV space, representing the distance of the center of mass of the entire **CQ**^0^ molecule to the center of the membrane (see Figure S12a in SI).

(4 1-palmitoyl-2-oleoyl-sn-glycero-3-phosphocholine

(5 1-palmitoyl-2-oleoyl-sn-glycero-3-phosphoserine

(6 We assume that the doubly charged **CQ^2+^** is not stable in the membrane, so that **CQ^2+^** is expected to be deprotonated in the aqueous phase to form the **CQ^+^** before entering the membrane. Under this assumption, the free energy surface would only be slightly shifted without an impact on the permeation mechanism.

