## Supporting Information for "Physical Chemistry of Drug Permeation through the Cell Membrane with Atomistic Detail"

### S1. $CQ^0$ Permeation through a POPC Membrane

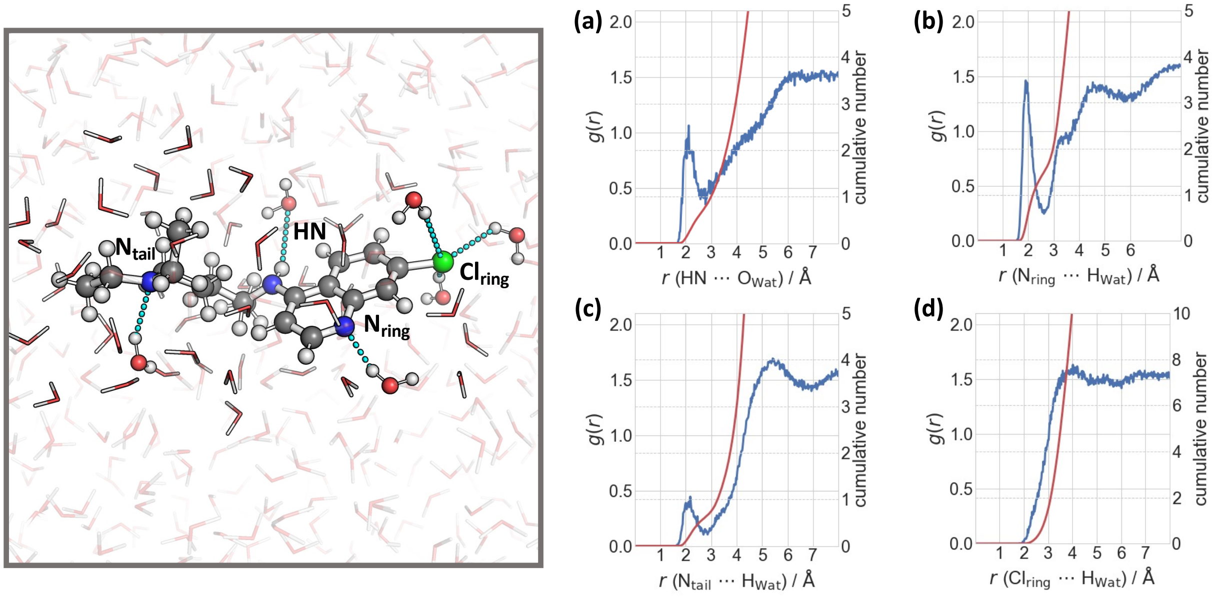


Figure S1: $\mathbf{CQ^0}$ in the solvent state. Left: Representative snapshot of $\mathbf{CQ^0}$ in the aqueous phase, labeling its H-bond acceptor and donor atoms. Dashed lines indicate H-bond interactions with water molecules. Right: Radial distribution functions and coordination numbers of H-bond interactions for (a) $HN\cdots O_{Wat}$, (b) $N_{ring}\cdots H_{Wat}$, (c) $N_{tail}\cdots H_{Wat}$, and (d) $Cl_{ring}\cdots H_{Wat}$ of $\mathbf{CQ^0}$ in the aqueous phase.

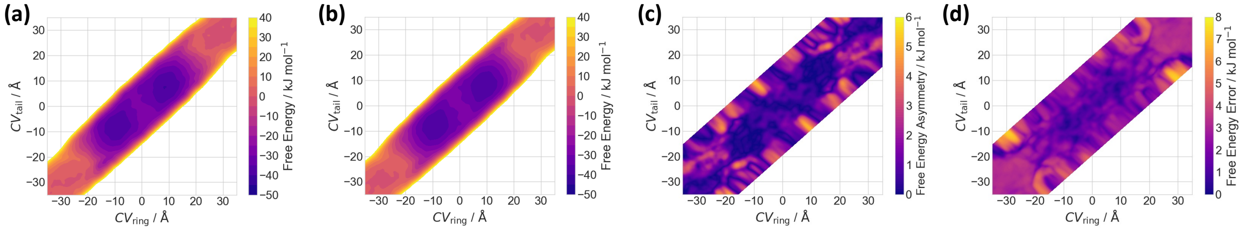


Figure S2: Analysis of the free energy surface for the permeation process of $\mathbf{CQ^0}$ through the POPC membrane. (a) Free energy surface as a function of $CV_{ring}$ and $CV_{tail}$ (see Figure 1 in the main text). (b) Symmetrized free energy surface: $G_{sym} = 1/2[G(CV_{ring}, CV_{tail}) + G(-CV_{ring}, -CV_{tail})]$. (c) Asymmetry analysis of the free energy surface: $G_{asym} = \mathrm{abs}[G(CV_{ring}, CV_{tail}) - G(-CV_{ring}, -CV_{tail})]$. (d) Estimated free energy error from block averaging of the symmetrized free energy profile.

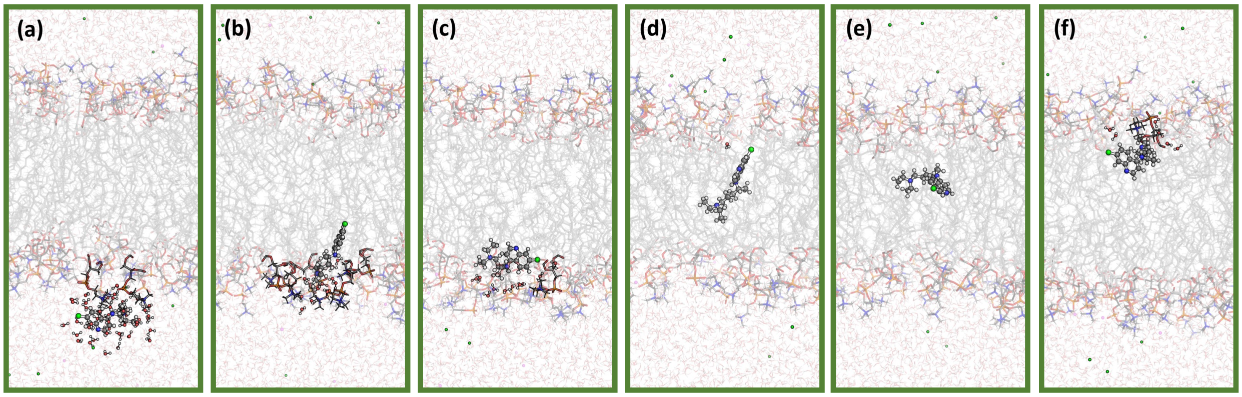


Figure S3: (a)–(f) Representative snapshots of the $\mathbf{CQ^0}$ permeation process across the POPC membrane.

### S2. $\mathbf{CQ^+}$ Permeation through a POPC Membrane

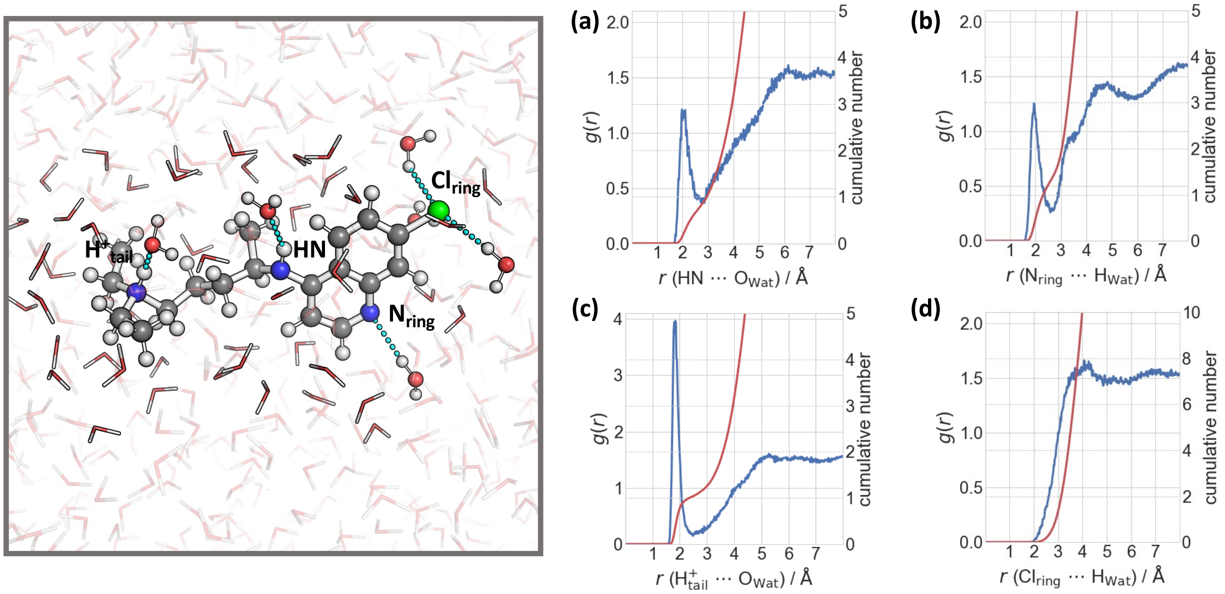


Figure S4: $\mathbf{CQ^+}$ in the solvent state. Left: Representative snapshot of $\mathbf{CQ^+}$ in the aqueous phase, labeling its H-bond acceptor and donor atoms. Dashed lines indicate H-bond interactions with water molecules. Right: Radial distribution functions and coordination numbers of H-bond interactions for (a) $\mathrm{HN}\cdots\mathrm{O_{Wat}}$, (b) $\mathrm{N_{ring}}\cdots\mathrm{H_{Wat}}$, (c) $\mathrm{H^+}_{\mathrm{tail}}\cdots\mathrm{O_{Wat}}$, and (d) $\mathrm{Cl_{ring}}\cdots\mathrm{H_{Wat}}$.

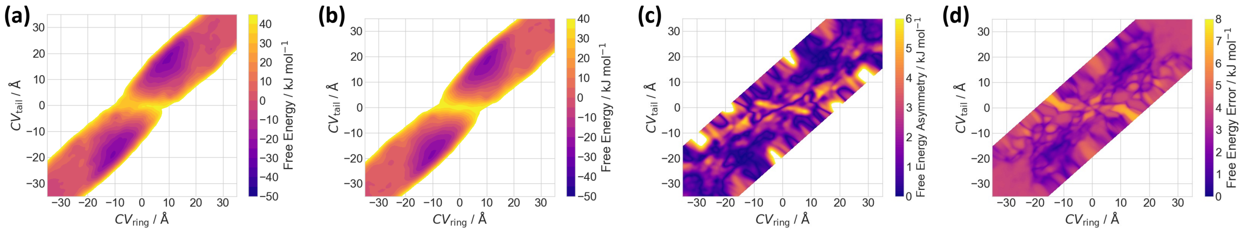


Figure S5: Analysis of the free energy surface for the permeation process of $\mathbf{CQ^+}$ through the POPC membrane. (a) Free energy surface as a function of $\mathrm{CV_{ring}}$ and $\mathrm{CV_{tail}}$ (see Figure 1 in the main text). (b) Symmetrized free energy surface: $G_{\mathrm{sym}} = 1/2[G(\mathrm{CV_{ring}}, \mathrm{CV_{tail}}) + G(-\mathrm{CV_{ring}}, -\mathrm{CV_{tail}})]$. (c) Asymmetry analysis of the free energy surface: $G_{\mathrm{asym}} = \mathrm{abs}[G(\mathrm{CV_{ring}}, \mathrm{CV_{tail}}) - G(-\mathrm{CV_{ring}}, -\mathrm{CV_{tail}})]$. (d) Estimated free energy error from block averaging of the symmetrized free energy profile.

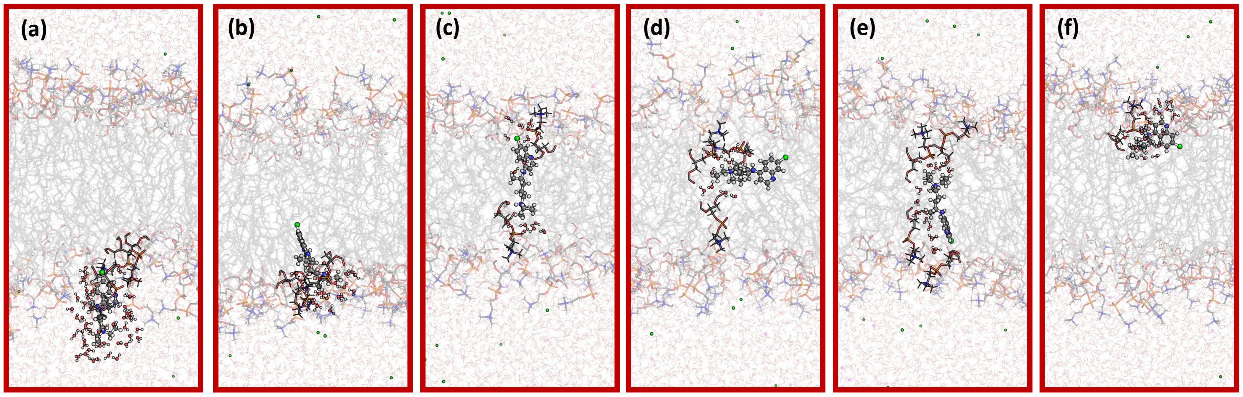


Figure S6: (a)–(f) Representative snapshots of the $\mathbf{CQ^+}$ permeation process across the POPC membrane.

#### S3. $\mathbf{CQ^0}\rightleftharpoons\mathbf{CQ^+}$ Permeation through a POPC Membrane

We used the thermodynamic cycle depicted in Figure S7a to calculate the free energy for the proton transfer, and its associated p$K_\mathrm{a}$ value, in the lipid phase.[S1] The transfer free energies from the aqueous to the lipid phase for $\mathbf{CQ^0}$ ($\Delta G_\mathrm{trans;neutral}$) and $\mathbf{CQ^+}$ ($\Delta G_\mathrm{trans;charged}$) were calculated by the well-tempered metadynamics simulations. The term $\Delta G_{\mathbf{W};\mathrm{H^+-transfer}}$ corresponds to the deprotonation free energy of $\mathbf{CQ^+}$ in the aqueous phase, which is calculated by

$$\Delta G_{\mathbf{W};\mathrm{H^+-transfer}} = \ln(10)RT(\mathrm{p}K_\mathrm{a} - \mathrm{pH}) \quad . \tag{1}$$

Here, $R$ and $T$ are the universal gas constant and temperature, respectively. The experimental p$K_\mathrm{a}$ of 10.4[S2] and a neutral pH (7) were used. The free energy difference between $\mathbf{CQ^+}$ and $\mathbf{CQ^0}$ then amounts to 20.2 kJ mol$^{-1}$ at 310 K. This value was used as a constant shift for the calculated free energy surface of $\mathbf{CQ^0}$.

Summing up the individual terms, the deprotonation free energy of $\mathbf{CQ^+}$ could be determined for the lipid phase ($\Delta G_{\mathbf{L};\mathrm{H^+-transfer}}$). The associated p$K_\mathrm{a}$ value is then given by

$$\mathrm{p}K_\mathrm{a} = \frac{\Delta G_{\mathbf{L};\mathrm{H^+-transfer}}}{\ln(10)RT} + \mathrm{pH} \quad . \tag{2}$$

Finally, the fraction of $\mathbf{CQ^+}$ in the different environments is given by

$$\frac{[\mathbf{CQ^+}]}{[\mathbf{CQ^0}] + [\mathbf{CQ^+}]} = \frac{\exp(-\beta G_{\mathbf{L};\mathrm{H^+-transfer}})}{1 + \exp(-\beta G_{\mathbf{L};\mathrm{H^+-transfer}})} \tag{3}$$

where $\beta$ is the reciprocal thermodynamic temperature $(RT)^{-1}$. This fraction of $\mathbf{CQ^+}$, and its complementary one for $\mathbf{CQ^0}$, were used to weight the free energy surfaces of $\mathbf{CQ^+}$ and $\mathbf{CQ^0}$ to determine the protonation-dependent $\mathbf{CQ^0} \rightleftharpoons \mathbf{CQ^+}$ free energy surface.

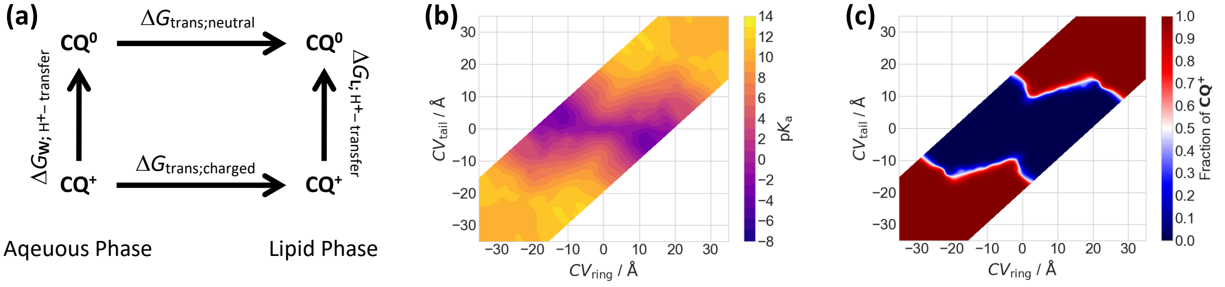


Figure S7: Analysis of the position-dependent equilibrium of $\mathbf{CQ^0}$ and $\mathbf{CQ^+}$ for the permeation through the POPC membrane. (a) Thermodynamic cycle for the calculation of the free energy difference $\Delta G_{\mathbf{L};\mathrm{H^+-transfer}}$ of $\mathbf{CQ^0}$ and $\mathbf{CQ^+}$ in the lipid phase. The deprotonation free energy $\Delta G_{\mathbf{W};\mathrm{H^+-transfer}}$ in the aqueous phase was calculated from Equation (1), while the two $\Delta G_{\mathrm{trans}}$ were calculated from well-tempered metadynamics simulations (see text above). (b) p$K_\mathrm{a}$ value of **CQ** as a function of $\mathrm{CV_{ring}}$ and $\mathrm{CV_{tail}}$ (see Figure 1 in the main text), calculated from Equation (2). (c) Fraction of $\mathbf{CQ^+}$ as a function of $\mathrm{CV_{ring}}$ and $\mathrm{CV_{tail}}$, calculated from Equation (3).

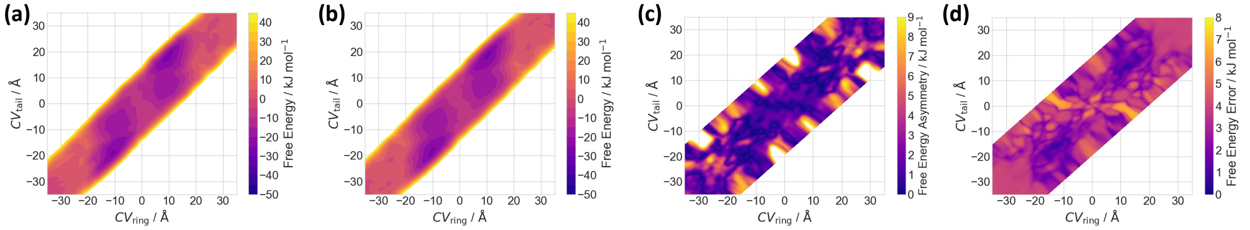


Figure S8: Analysis of the free energy surface for the permeation process of $\mathbf{CQ^0} \rightleftharpoons \mathbf{CQ^+}$ through the POPC membrane. (a) Free energy surface as a function of $\mathrm{CV_{ring}}$ and $\mathrm{CV_{tail}}$ (see Figure 1 in the main text). (b) Symmetrized free energy surface: $G_{\mathrm{sym}} = 1/2[G(\mathrm{CV_{ring}}, \mathrm{CV_{tail}}) + G(-\mathrm{CV_{ring}}, -\mathrm{CV_{tail}})]$. (c) Asymmetry analysis of the free energy surface: $G_{\mathrm{asym}} = \mathrm{abs}[G(\mathrm{CV_{ring}}, \mathrm{CV_{tail}}) - G(-\mathrm{CV_{ring}}, -\mathrm{CV_{tail}})]$. (d) Estimated free energy error from block averaging of the symmetrized free energy profile.

### S4. $\mathbf{CQ^0}$ Permeation through a Mixed POPC/POPS Membrane

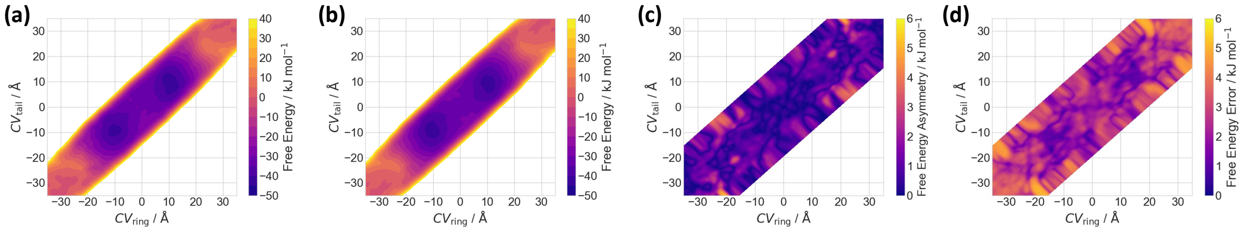


Figure S9: Analysis of the free energy surface for the permeation process of $\mathbf{CQ^0}$ through the POPC/POPS membrane. (a) Free energy surface as a function of $\mathrm{CV_{ring}}$ and $\mathrm{CV_{tail}}$ (see Figure 1 in the main text). (b) Symmetrized free energy surface: $G_{\mathrm{sym}} = 1/2[G(\mathrm{CV_{ring}}, \mathrm{CV_{tail}}) + G(-\mathrm{CV_{ring}}, -\mathrm{CV_{tail}})]$. (c) Asymmetry analysis of the free energy surface: $G_{\mathrm{asym}} = \mathrm{abs}[G(\mathrm{CV_{ring}}, \mathrm{CV_{tail}}) - G(-\mathrm{CV_{ring}}, -\mathrm{CV_{tail}})]$. (d) Estimated free energy error from block averaging of the symmetrized free energy profile.

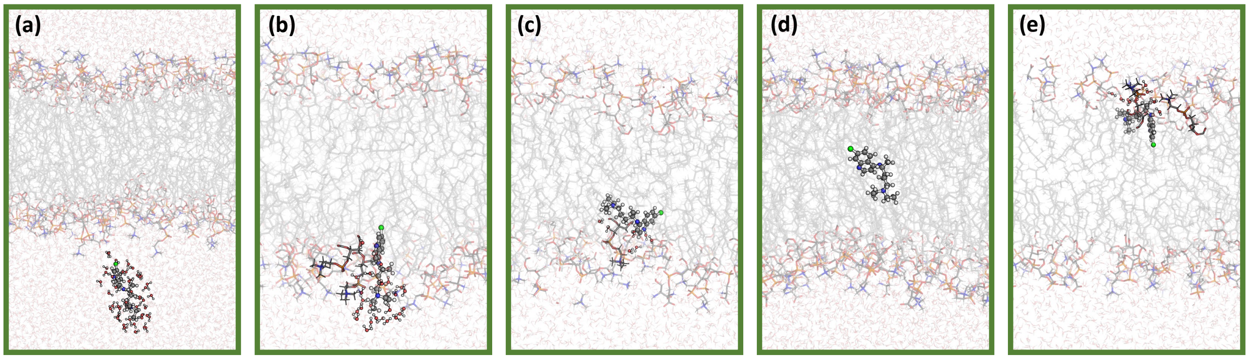


Figure S10: (a)–(f) Representative snapshots of the $\mathbf{CQ^0}$ permeation process across the POPC/POPS membrane.

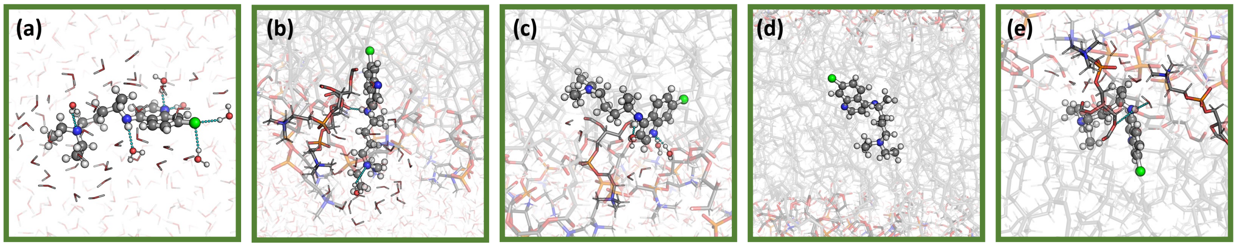


Figure S11: (a)–(e): Representative snapshots of the $\mathbf{CQ^0}$ permeation process across the POPC/POPS membrane, showing the interactions of $\mathbf{CQ^0}$ with water molecules and POPC/POPS headgroups.

### S5. $CQ^0$ Permeation through a POPC Membrane with an External Field

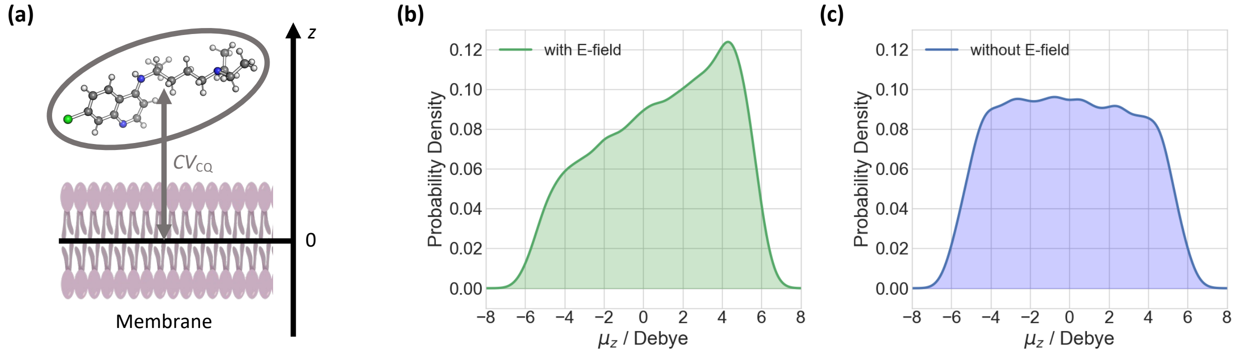


Figure S12: (a) Schematic representation of the employed CV to determine the impact of an external electric field. (b) Distribution of the $z$-component of the dipole moment ($\mu_Z$) of $\mathbf{CQ^0}$ from well-tempered metadynamics simulations with an external electric field of 10 mV $\text{Å}^{-1}$. (c) Distribution of the $z$-component of the dipole moment ($\mu_Z$) of $\mathbf{CQ^0}$ from well-tempered metadynamics simulations without an external electric field.

### S6. Calculation of Properties

**Diffusion Coefficients.** The position-dependent diffusion coefficients $D(z_i)$ of the drugs can be derived from the solution of the generalized Langevin equation (GLE), in which the drug is considered as a harmonic oscillator in a frictional bath.[S3,S4] The diffusion coefficient of the drug reads:[S3,S4]

$$D(z_i) = \lim_{s\to 0} \frac{-\mathcal{L}\{C_v(t)\}(s) \cdot \langle \delta z^2 \rangle_i \langle \dot{z}^2 \rangle_i}{\mathcal{L}\{C_v(t)\}(s) \cdot [s\langle \delta z^2 \rangle_i + s^{-1}\langle \dot{z}^2 \rangle_i] - \langle \delta z^2 \rangle_i \langle \dot{z}^2 \rangle_i} \quad (4)$$

where $z$ corresponds to the distance of the molecule to the membrane center, as defined by the collective variable (see Figure 1 in the main text). $\delta z$ is the displacement from its average position, $\langle z^2 \rangle$ and $\langle \dot{z}^2 \rangle$ are the variances of the position and the velocity, respectively. The velocity autocorrelation function $C_v(t)$ is numerically calculated as

$$C_v(t) = \langle \dot{z}(0)\dot{z}(t) \rangle = \frac{1}{n} \sum_{j=0}^{n} \dot{z}(j \cdot \Delta t)\dot{z}(t + j \cdot \Delta t) \tag{5}$$

where $n$ is the number of sampled points and $\Delta t$ the time step. Its Laplace transform $\mathcal{L}\{C_v(t)\}(s)$ is given by

$$\mathcal{L}\{C_v(t)\}(s) = \int_0^\infty C_v(t) \exp(st) \mathrm{d}t \tag{6}$$

which is a function of the transformation parameter $s$.

Equation (4) was solved numerically following the protocol in Ref. S5. The drugs were restrained using a harmonic potential along the $z$-axis (see Figure S12a). The drugs were then pulled through the membrane to the opposite site (at +40 Å) using a moving harmonic restraint with a force constant of 100 kJ mol$^{-1}$ Å$^{-2}$ in 80 ns *NPT* simulations. The procedure was carried out for 3 replicas for $\mathbf{CQ^0}$ and $\mathbf{CQ^+}$, considering the POPC membrane, as well as for $\mathbf{CQ^0}$ across the mixed POPC/POPS membrane.

In a first set of calculations, we determined the full diffusion coefficient profile of $\mathbf{CQ^0}$ across the POPC membrane. From the pulling simulations, snapshots were taken in 1 ns intervals. This leads to 81 windows in 1 Å steps in the range of $z$ = [–40 Å; +40 Å]. For each window $i$, the system was re-equilibrated for 10 ns *NPT* simulation by restraining the position of $\mathbf{CQ^0}$ at $z_i$ with a harmonic potential of force constant 125 kJ mol$^{-1}$ Å$^{-2}$, following 5 ns production time in the *NVE* ensemble. The MD data of each of the 81 windows were numerically processed according to Equations (4–6) for $s$ ranging from 0.005 to 0.1 fs$^{-1}$, following a procedure similar to that of Ref. S5. An almost linear segment was defined by a curvature of the diffusion coefficient (as a function of the parameter $s$) lower than 0.01 Å$^2$ fs. The position-dependent diffusion coefficient $D(z_i)$ was then determined by the intercept of a linear regression from this segment. The final diffusion profile was obtained by averaging over the replicas and smoothening with a moving average using five windows. The results are shown in Figure S13.

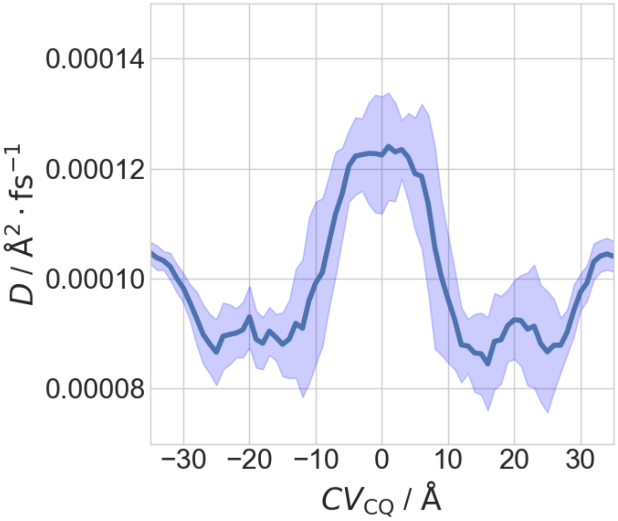


Figure S13: Position-dependent diffusion coefficient of CQ0 upon permeation through a POPC membrane.

In a second set of calculations, we determined the local diffusion coefficients along the minimum free energy path (MFEP) within our two dimensional CV space (see Figure 1 in the main text) for all systems. We followed the procedure described in Refs. S6,S7. Here, we only considered the positions at the free energy maxima, because these contribute most to the permeability of the drugs, as demonstrated in Refs. S6,S7, thereby significantly reducing the computational cost. We applied the same computational protocol as described above, except that a two dimensional harmonic potential was used to restrain the ring and tail moieties of the drugs ($CV_{ring}$ and $CV_{tail}$, respectively). The deviations from the reference positions were projected onto the tangent of the MFEP to obtain a time series for the diffusion along permeation coordinate.[S6,S7] The latter was then processed to determine the diffusion coefficient, using the protocol as described above. The results are summarized in Table S1.

Table S1: Position-dependent diffusion coefficients of $\mathbf{CQ^0}$ and $\mathbf{CQ^+}$, considering the POPC and POPC/POPS membranes.

| | System | $CV_{ring}$ / Å | $CV_{tail}$ / Å | $D$ / $10^4$ Å$^2$ fs$^{-1}$ |
|---|---|---|---|---|
| $\mathbf{CQ^0}$ | POPC | −22.69 | −22.15 | $2.10 \pm 0.16$ |
| $\mathbf{CQ^0}$ | POPC/POPS | −22.59 | −29.12 | $1.32 \pm 0.02$ |
| $\mathbf{CQ^+}$ | POPC | 3.30 | −0.73 | $1.63 \pm 0.14$ |
| $\mathbf{CQ^+}$ | POPC | −23.30 | −24.80 | $1.51 \pm 0.19$ |

**Permeability Coefficients.** The permeability coefficient $P$ was calculated from the inhomogeneous solubility-diffusion model by[S8]

$$\frac{1}{P} = \exp(-\beta G_{\mathrm{ref}}) \int_{z_1}^{z_2} \frac{\exp(\beta G(z))}{D(z)} \mathrm{d}z \tag{7}$$

where $z$ corresponds to the permeation coordinate. $G(z)$ and $D(z)$ are the position-dependent free energy and diffusion coefficient, respectively, and $\beta$ is the reciprocal thermodynamic temperature. $G_{\mathrm{ref}}$ is the free energy in the aqueous phase, which was appropriately shifted to be zero.

The integral was calculated along the minimum free energy path (MFEP), using the diffusion coefficients reported in Table S1. The integration boundaries $z_{1/2}$ correspond to the positions of **CQ** at $CV_{ring} = \pm 35$ Å and $CV_{tail} = \pm 35$ Å. For the simulation of $\mathbf{CQ^0}$ across the POPC membrane using the one-dimensional $CV_{CQ}$, the position-dependent diffusion profile was used (see Figure S12). The integration boundaries were set to $z_{1/2} = \pm 35$ Å. The integrations were done numerically, using the trapezoidal rule. The error was estimated by adding Gaussian distributed noise to the integrand, according to its mean value and standard deviation at each integration point. We integrated $10^7$ sampled profiles to calculate the standard deviation of the permeability coefficient. The results are shown in Table S2.

Table S2: Permeability coefficients of **CQ**, considering the POPC and POPC/POPS membranes.

| System | | | $P$ / cm s$^{-1}$ |
|---|---|---|---|
| $\mathbf{CQ^0}$ | POPC | $CV_{CQ}$ | $27.6 \pm 9.0$ |
| $\mathbf{CQ^0}$ | POPC | $(CV_{ring},CV_{tail})$ | $38.2 \pm 7.8$ |
| $\mathbf{CQ^+}$ | POPC | $(CV_{ring},CV_{tail})$ | $8.5\ 10^{-4} \pm 8.5\ 10^{-4}$ |
| $\mathbf{CQ^0} \rightleftharpoons \mathbf{CQ^+}$ | POPC | $(CV_{ring},CV_{tail})$ | $26.0 \pm 6.0$ |
| $\mathbf{CQ^0}$ | POPC/POPS | $(CV_{ring},CV_{tail})$ | $44 \pm 12$ |
